# Switch-like Compaction of Poly(ADP-ribose) Upon Cation Binding

**DOI:** 10.1101/2023.03.11.531013

**Authors:** Mohsen Badiee, Adam L. Kenet, Laura R. Ganser, Tapas Paul, Sua Myong, Anthony K. L. Leung

## Abstract

Poly(ADP-ribose) (PAR) is a homopolymer of adenosine diphosphate ribose that is added to proteins as a post-translational modification to regulate numerous cellular processes. PAR also serves as a scaffold for protein binding in macromolecular complexes, including biomolecular condensates. It remains unclear how PAR achieves specific molecular recognition. Here, we use single-molecule fluorescence resonance energy transfer (smFRET) to evaluate PAR flexibility under different cation conditions. We demonstrate that, compared to RNA and DNA, PAR has a longer persistence length and undergoes a sharper transition from extended to compact states in physiologically relevant concentrations of various cations (Na^+^, Mg^2+^, Ca^2+^, and spermine). We show that the degree of PAR compaction depends on the concentration and valency of cations. Furthermore, the intrinsically disordered protein FUS also served as a macromolecular cation to compact PAR. Taken together, our study reveals the inherent stiffness of PAR molecules, which undergo switch-like compaction in response to cation binding. This study indicates that a cationic environment may drive recognition specificity of PAR.

**Significance:** Poly(ADP-ribose) (PAR) is an RNA-like homopolymer that regulates DNA repair, RNA metabolism, and biomolecular condensate formation. Dysregulation of PAR results in cancer and neurodegeneration. Although discovered in 1963, fundamental properties of this therapeutically important polymer remain largely unknown. Biophysical and structural analyses of PAR have been exceptionally challenging due to the dynamic and repetitive nature. Here, we present the first single-molecule biophysical characterization of PAR. We show that PAR is stiffer than DNA and RNA per unit length. Unlike DNA and RNA which undergoes gradual compaction, PAR exhibits an abrupt switch-like bending as a function of salt concentration and by protein binding. Our findings points to unique physical properties of PAR that may drive recognition specificity for its function.

## Introduction

Poly(ADP-ribose) (PAR) is a homopolymer of adenosine diphosphate (ADP)-ribose and a post-translational modification that regulates DNA repair, transcription, translation and the cell cycle (1–4). PAR homopolymers were recently proposed to serve as multivalent scaffolds that mediate the formation of subcellular structures, such as stress granules, nucleolus, and other biomolecular condensates, through non-covalent interactions (2). For example, PAR forms biomolecular condensates by recruiting DNA repair proteins at DNA damage sites upon genotoxic stress or RNA-binding proteins in stress granules upon translation inhibition (5, 6). Given that these cellular processes are tightly regulated and involve a selected set of PAR-binding proteins, PAR polymers likely serve as specific molecular recognition elements. However, how specificity is achieved in the context of a highly-charged and relatively simple homopolymer of repeating ADP-ribose units remains unknown.

One of the main challenges towards a more granular view of PAR’s specific recognition has been the lack of structural information. Although PAR exists as a homopolymer composed of 2–200 ADP-ribose units, currently available structures are limited to PAR dimers (7–9). Bulk level solution studies of longer PAR homopolymers using nuclear magnetic resonance (NMR) spectroscopy (Schultheisz et al., 2009) and circular dichroism (CD) (10) did not reveal the presence of a well-defined structure, although they showed that PAR structure may be sensitive to metal cations, multivalent cations such as spermine, and different salt (NaCl) concentrations. Notably, the CD analysis also showed that changes in environment surrounding the adenine bases of PAR detected at high salt concentrations were lost upon heating (>55 °C), suggesting a melting transition indicative of macromolecular structure. Therefore, these early studies have hinted at the possibility that PAR adopts distinct structures in a cation-dependent manner. However, biochemical, biophysical and structural analyses of PAR homopolymers is challenging given the dynamic and repetitive nature of these molecules, leading to very little progress being made in this area over the last few decades.

Single-molecule technology has been instrumental for evaluating structural and dynamic properties of nucleic acids, such as the flexibility of single-stranded DNA and RNA (11–13). However, single-molecule analyses have not been possible for PAR partly due to the lack of technology to label one or both of its termini. Here, using novel labeling techniques recently established by our team (14, 15), we developed a series of probes for single-molecule fluorescence resonance energy transfer (smFRET) studies of PAR homopolymers of defined lengths. We employed these probes to assess PAR flexibility, compare them with DNA and RNA, and evaluate the impact of biologically relevant cations on this fundamental property. We observed cation-dependent compaction of PAR.

## Results

### Development of smFRET probes for PAR flexibility

We designed the smFRET PAR probe following a well-established configuration (11), where a Cy5-labeled and biotin-conjugated single-stranded DNA is annealed to a chimeric strand composed of a complementary DNA sequence linked to a Cy3-labeled PAR (Fig. 1A). We used a combination of chemical and enzymatic methods to create the chimera ssDNA-PAR-Cy3 (Fig. S1A). Briefly, PAR generated from the synthetic enzyme PARP5a was cleaved by base hydrolysis, followed by HPLC purification to separate PAR chains of defined lengths, leaving a phosphate group at the 1” terminus (Fig. S1B-D). The phosphate was then functionalized with an alkyne-amine linker, which allows for conjugation with azide-labeled ssDNA (15). Cy3 dye was incorporated at the 2’ terminus of the alkyne-PAR using the enzymatic labeling method ELTA (14). Various lengths of ssDNA-PAR-Cy3 chimera were successfully generated (Fig. S1E-F; Methods), with an overall yield of 10-25%. Finally, the ssDNA-PAR-Cy3 chimeras were annealed to a complementary, biotinylated 18 mer-Cy5-DNA to form a partial duplex structure (pdPAR; Fig. 1A-B, S1A, S1G). This arrangement allows us to probe the end-to-end distance of various PAR lengths by smFRET in the same way used for measuring single-stranded DNA flexibility (11).

**Figure 1.**
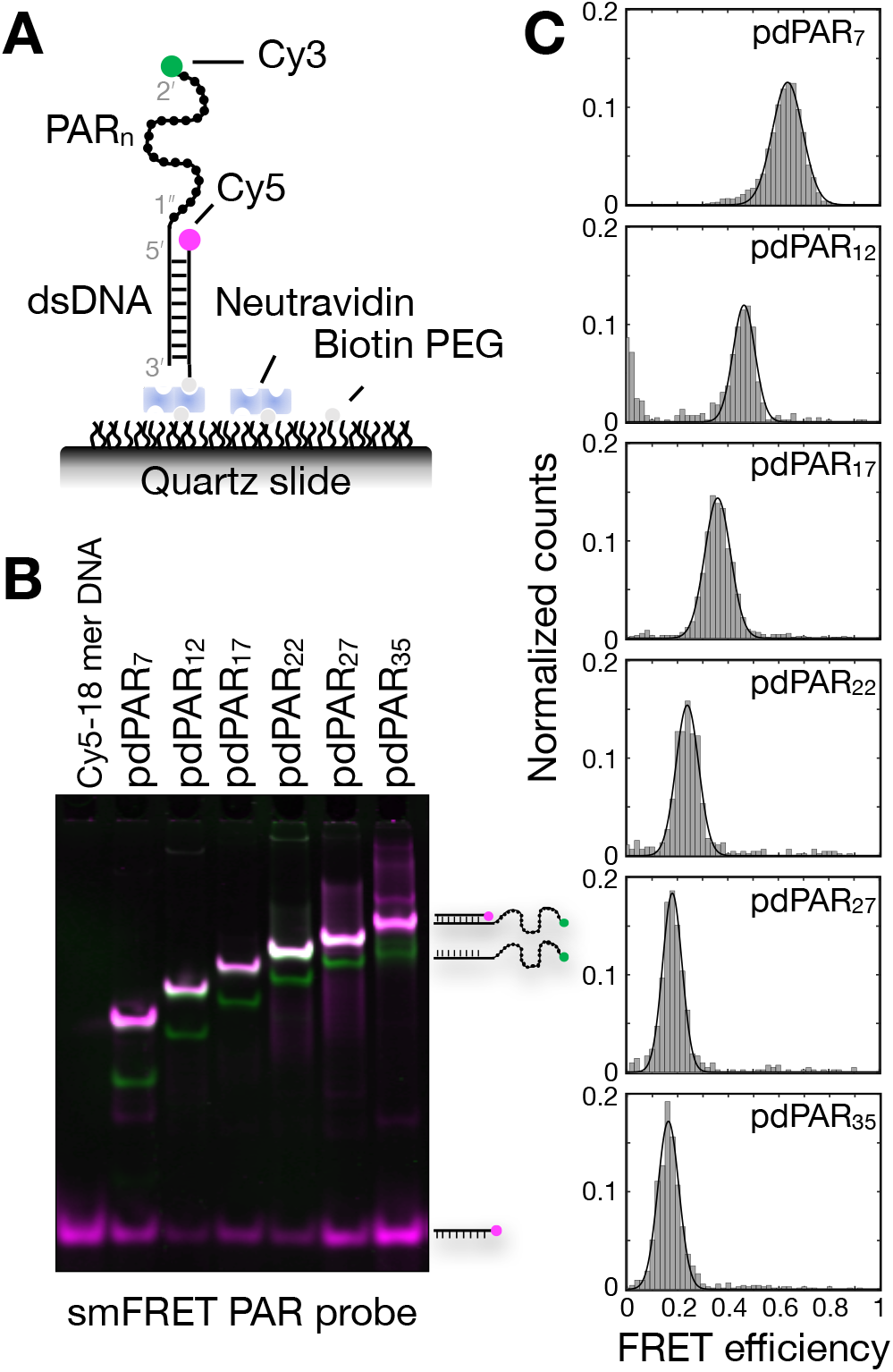
Development of smFRET probe to measure PAR flexibility. **A.** Schematic showing the configuration of pdPARn probe attached to single molecule surface. Cy3 attached to 2’ end of PAR and DNA attached to 1” end of PAR. Cy5 and biotin are on 5’ and 3’ end of complementary DNA strand, respectively. **B.** 15% polyacrylamide native gel analysis of pdPAR_n_ probes (n= 7, 12, 22, 27, 35), indicated with Cy3 (green) and Cy5 (magenta) signals. **C.** FRET histograms of pdPAR_7-35_ at 200 mM NaCl. Single-peak Gaussian distributions are used to fit the histograms after removing the donor-only population. The fit is shown as a solid black line.

To evaluate PAR flexibility, we made six smFRET probes (pdPAR_n_) to cover a range of chain lengths (n = 7, 12, 17, 22, 27, 35), which are expected to have an inverse relationship with the FRET efficiency. The formation of pdPAR_n_ was confirmed by native gel electrophoresis, which clearly distinguished the partial duplex from the unannealed single strands (Fig. 1B and S1G). We immobilized the FRET probes on a PEG passivated quartz slide (16) and acquired FRET values at 200 mM NaCl concentration (Fig. 1C). Cy3 and Cy5 intensities from 100-300 single molecules of pdPAR_n_ per field of view were collected upon excitation by a 532 nm laser, and FRET efficiency was calculated. FRET histograms were built by combining FRET values of over 2,000 single molecules collected from ~20 fields of view. Similar to single-stranded DNA, one major FRET peak was observed for each PAR probe, likely indicating a time-averaged value resulting from conformational fluctuations of individual PAR molecules that occur at a faster timescale than the sampling frequency (~100 ms). As expected, FRET efficiency (***E***) has an inverse relationship with the PAR length; *i.e*., longer PARs display lower FRET values. The FRET efficiency values, measured as the center of the fitted Gaussian peak, reflect the average end-to-end distance measurements and provide a suitable proxy for evaluating PAR polymer flexibility. Taken together, we developed a set of smFRET probes, pdPARs, and validated that these probes can be used to measure flexibility of PAR homopolymers of varying lengths.

### PAR is stiffer than single-stranded DNA and RNA at physiological salt condition

Next, we acquired FRET histograms for the six PAR lengths (7–35 mer) across a wide range of salt (NaCl) concentrations of 25–2000 mM (Table S1A). To deduce the polymer properties of PAR, we fitted the experimental FRET efficiency values into the worm-like chain (WLC) model (Fig. 2A), which has been used to describe the flexibility of single-stranded DNA and RNA (11, 13). The WLC model uses two parameters to define the mechanical properties of a polymer: contour length and persistence length. Contour length (L) is the length of a fully stretched polymer, and for a polymer with N monomers, *L* = *N*·*b_0_*, where *b_0_* is equivalent to the monomer length. We first deduced *b_0_* of PAR from 10 published ADP-ribose- or PAR-bound structures by measuring the average distance for four candidate monomeric units: ADP-ribose (11.6 Å), iso-ADP-ribose (11.5 Å), and, as measured for DNA and RNA, the distance between adjacent phosphates (□: 12.5 Å and β: 12.7 Å) (Fig. 2B and S2A-C). To compare our FRET data to the WLC model, we performed Metropolis Monte-Carlo simulations of the end-to-end distance distribution over time to calculate predicted FRET values (Fig. 2C and S2D; (11)). The average root mean square deviations (RMSD) were calculated between our experimental data and simulated values across various lengths and salt concentrations. The minimal RMSD was achieved when *b_0_* = 11.5-11.6 Å (Fig. 2B and S2E-F), which is consistent with the length of a single ADP-ribose unit calculated from the crystal structures.

**Figure 2.**
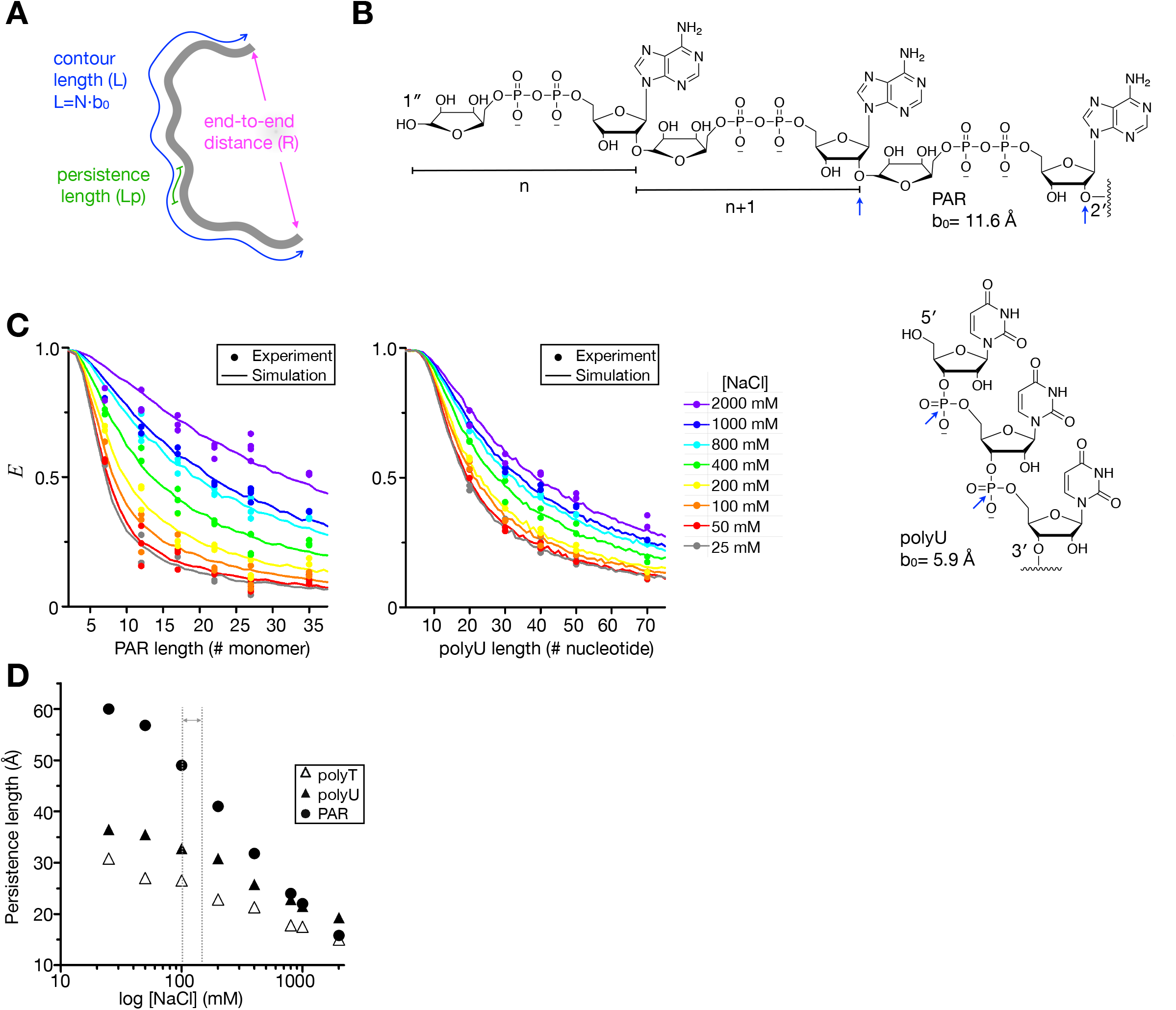
Worm-like chain model revealed PAR as stiffer than single-stranded DNA and RNA at physiological NaCl concentrations. **A.** Parameters of worm-like chain (WLC) model are illustrated. **B.** Chemical structures of linear 3 mer-PAR and 3 nt-polyU. The blue arrows indicates how monomeric size (b0) are defined. Please note that the monomeric unit of PAR is twice the size of RNA. **C.** Simulated (***E***) and experimental (***E***_FRET_) energy transfer values over various lengths of PAR and polyU were plotted for eight tested NaCl concentrations from 2 M NaCl (violet) to 25 mM (gray), with the corresponding persistence lengths at each concentration indicated. **D.** Comparison of persistence length between PAR, ssRNA and ssDNA. Data from this study and published values for polyT (Ref.11) were plotted across NaCl concentration in log scale. The physiological concentration range of NaCl (100-150 mM) are highlighted.

The other parameter used to describe polymer flexibility is persistence length (Lp). Persistence length represents the length scale over which a polymer chain maintains a certain direction. The longer the persistence length, the stiffer the polymer is, or, in other words, the more energy is required to bend it (17). With *b_0_* defined as the monomeric ADP-ribose (11.6 Å) in the WLC model, the persistence length of PAR is determined to be 60.0 Å at 25 mM NaCl, which is significantly longer than oligo dT DNA (30.8 Å; Fig. 2D and S2G). Lp of charged polymers is known to depend on the electrostatic repulsion within the backbone and the presence of counterions (i.e., charge screening), which reduces the bending stiffness and hence its persistence length (18–20). Thus, we measured Lp values at 2 M NaCl and observed a reduction in the persistence length (15.8 Å), which is comparable to DNA (15.0 Å). Altogether, these data indicate that the longer persistence length observed for PAR at low salt is mainly driven by the electrostatic repulsion of negative charges within the polymer, especially considering there are two phosphates per unit of PAR compared to one in DNA and RNA.

Since PAR is composed of ribose rather than deoxyribose, we compared the PAR homopolymers to RNA. Although PAR is composed of adenosines, we used polyU instead of polyA because the latter has the propensity to form a helical structure (21), which is not observed in PAR under physiological condition (22). We made a similar smFRET probe for RNA (23), where an 18 mer Cy5-labeled and biotinylated single-stranded RNA is annealed to a complementary RNA with a defined length of Cy3-labeled polyU. We then measured the FRET efficiency for a range of RNA lengths (20, 30, 40, 50, 70 mers) across the salt concentrations of 25–2000 mM (Table S1A; Fig. 2C and S2H). We also deduced the *b_0_* of RNA as 5.9 Å from 11 polyU-bound protein structures (Fig. S2C). Based on the WLC model with *b_0_* set at 5.9 Å, the persistence lengths of RNA were best described as 37 Å at 25 mM NaCl and 19 Å at 2 M NaCl (Fig. 2C and S2D-G), which are slightly higher than the published values for a freely diffused U_40_ construct (13). This expected difference can be attributed to the sensitivity of nucleic acid conformation to its local environment, where the presence of an 18-bp duplex in our FRET construct can cause the RNA to appear more extended, as shown previously (13). Notably, PAR has the longest persistence length compared to DNA and RNA at physiological NaCl concentrations (100-150 mM NaCl; Fig. 2D). Taken together, our analysis revealed that PAR is the stiffest amongst these nucleic acids.

### PAR compaction increases with salt valency

Metal cations, such as sodium and magnesium, play an essential role in folding, molecular recognition, and catalytic activity of RNA and DNA through various mechanisms, including reducing the repulsion between negative charges on the phosphate backbone (24, 25). Because a PAR monomer has one extra phospho-ribose moiety relative to each RNA/DNA monomer, it is expected that cations influence the charge screening, and, thus, compaction properties of PAR differently. Indeed, among these three nucleic acids, PAR exhibited the largest variation in persistence length (15.8-60.0 Å) across the NaCl concentrations tested compared with DNA (15.0-30.8 Å) and RNA (19.3-36.5 Å) (Fig. 2D and S2G). Therefore, PAR undergoes a higher degree of compaction by the monovalent cation (Na^+^) than DNA and RNA. To test the effect of divalent cations on PAR compaction, we applied Mg^2+^ and Ca^2+^, two cations with well-established roles in DNA, RNA (26–28) and PAR biology (25, 29, 30). The 27 mer PAR probe was chosen for subsequent analyses based on the large dynamic range of FRET values exhibited for the NaCl titrations (Table S1A). As in the case of Na^+^, the FRET efficiency increased as a function of Mg^2+^ and Ca^2+^ concentrations (0.5 – 50 mM), appearing as one major FRET peak per condition (Fig. S3). To evaluate the charge screening of cations as a function of valency, we plotted the persistence length over the salt concentrations (Fig. 3A), fitted to 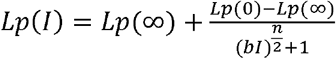 as in (13), and determined ***C***_**mid**_, *i.e*., the cation concentration required for the polymer to reach the halfway point between the most extended and the most compact states (Fig. 3B and Table S2). We observed that the concentration of divalent Mg^2+^ or Ca^2+^ required to reach the same compaction of PAR by monovalent Na^+^ was over two orders of magnitude lower. These data indicate that cations with higher valency screen negative charges significantly more effective than the monovalent cation.

**Figure 3.**
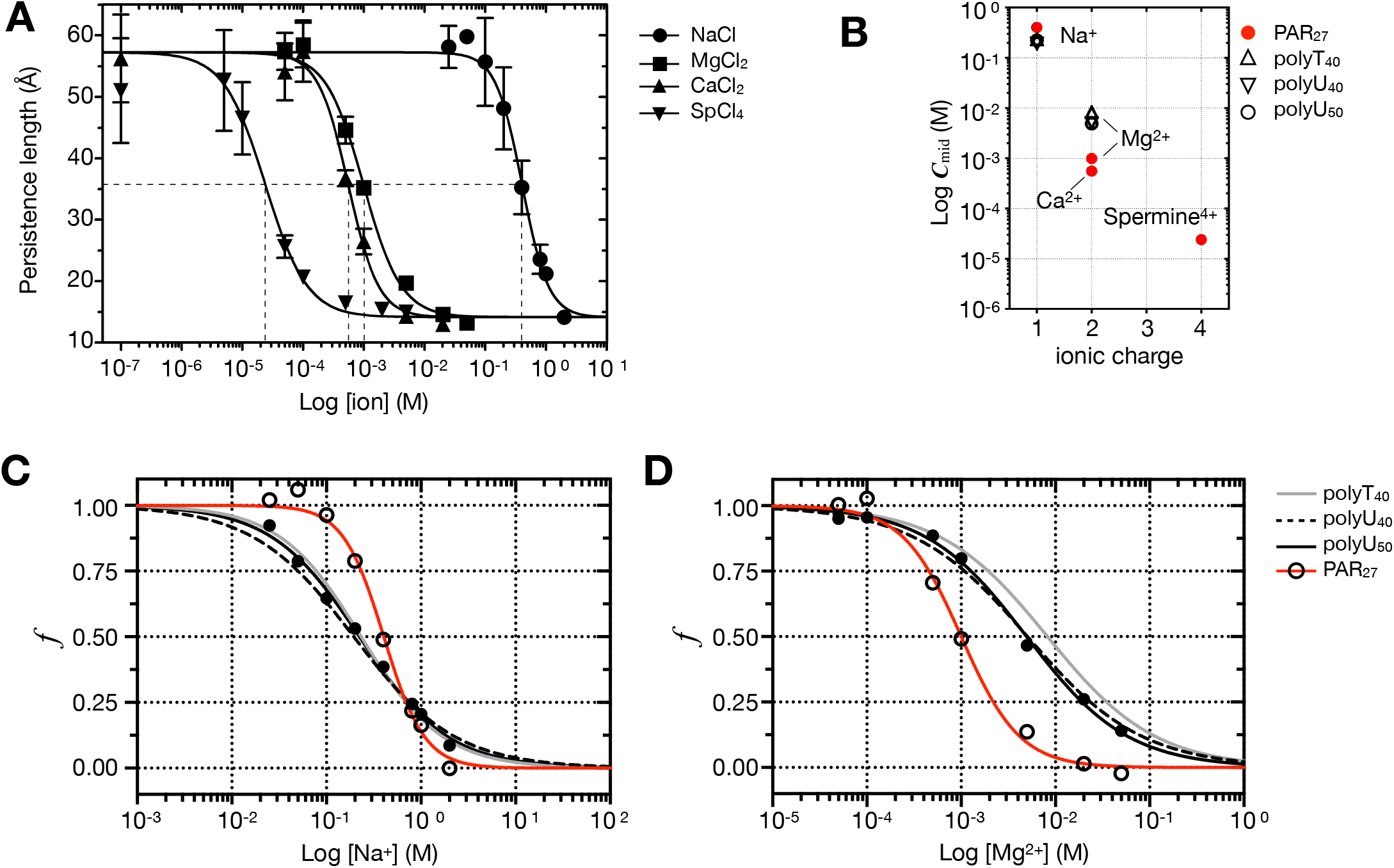
PAR is compacted by metals and spermine in a valency-dependent manner. **A.** Persistence lengths of pdPAR_27_ are plotted over cation concentrations in log scale. The data was fitted with a power-law equation (See text for equation). The intersection lines indicate ***c***mid of respective cations. **B.** The concentration of cations, ***c***mid, was extracted at mid point of Lp from panel A. The log scale of ***c***mid were plotted over the positive charge of the cation for PAR, polyU, and polyT. **C.** The fractions of persistence length ***f*** for PAR_27_, polyU_40_, polyU_50_ and poly T_40_ were plotted over NaCl concentrations. **D.** The fractions of persistence length ***f*** for PAR_27_, polyU_40_, polyU_50_ and poly T_40_ were plotted over MgCl_2_ concentrations. *Published values of polyU_40_ and poly T_40_ (Ref.13) were used as comparison in panels B-D.

Next, we tested the effect of spermine, a tetravalent polyamine cation known to condense DNA and some RNAs (31), and activate PAR synthesis (30). Raising the concentration of tetravalent spermine from 0.05 to 5 mM resulted in an increase in FRET efficiency of pdPAR_27_ probe (Fig. S3). The pattern of increase was similar to what we observed when using other cations, although the effects were achieved at significantly lower concentrations of spermine. Notably, the ***C***_**mid**_ for spermine was two orders of magnitude lower than that for divalent Mg^2+^/Ca^2^ and four orders of magnitude lower than that of the monovalent Na^+^(Fig. 3A and B). These data clearly indicate that PAR compaction strongly depends on the valency of cations. This phenomenon is consistent with other polyelectrolytes (32, 33), where multivalent ions enhance screening efficiency.

### PAR compacts sharply compared with DNA and RNA

Next, we compared PAR_27_ with other nucleic acids of similar length, utilizing both our own smFRET analyses on U_50_ (Table S1B) and published studies on U_40_/T_40_ across various Na^+^ or Mg^2+^ concentrations (13). Given that each nucleic acid has different persistence lengths representing their most extended [Lp(0)] and most compact [Lp(∞)] states, we plotted all data as scaled persistence length 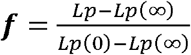 and determined the ***C***_**mid**_ as the cation concentration at which ***f*** = 0.5. The persistence length of each nucleic acid at two extremes of salt concentration represented as Lp(0) and Lp(∞) in the equation, and these two limiting quantities were extracted after fitting from plotting Lp over concentration (Fig. 3A). The ***C***_**mid**_ for PAR was higher than DNA and RNA for Na^+^, suggesting that a higher concentration of Na^+^ is required to compact PAR (Fig. 3C and Table S2). This difference likely arises from having to neutralize two phosphate groups for PAR monomer compared with one in RNA and DNA to achieve the same degree of compaction. In contrast, PAR required ~ 5–8 fold less Mg^2+^ for compaction than the other two nucleic acids (Fig. 3D and Table S2), which suggests that Mg^2+^ more effectively screens tandem phosphates in PAR backbone than single phosphates in DNA and RNA.

In addition to the different requirements of Na^+^ and Mg^2+^ for the compaction of PAR relative to DNA and RNA, we also observed a distinct pattern in nucleic acids’ response to the change in cation concentrations (Fig. 3C and D). We observed steeper slopes (***n***, Table S2) for PAR than for DNA and RNA. Within a narrow range of cation concentrations, PAR is significantly more responsive to the change in cation concentrations than DNA and RNA irrespective of the cation identity. These results indicate that PAR compacts via a distinct mechanism whereby PAR homopolymer resists compaction until cation concentration reaches a threshold at which PAR undergoes a large degree of compaction. This switch-like compaction, which we observed under physiological Mg^2+^ concentrations, may have implications for PAR-dependent cellular mechanisms.

### Switch-like PAR compaction is induced by FUS

In addition to small molecules and metal cations, PAR also interacts with numerous macromolecules, such as proteins, when serving as a multivalent scaffold. However, whether these interactions impact PAR flexibility has not been explored. Our recent study showed that PAR induces condensation of FUS protein via formation of transient interactions that “seed” the condensation process (34). FUS protein is a mostly disordered RNA binding protein (pI = 9.4), featuring multiple positively charged domains (35). To explore the binding between PAR and FUS further, we tested whether PAR can be compacted by FUS in the range of 5–500 nM. Interestingly, PAR was compacted by FUS in a switch-like manner as seen in the small biomolecule cations (Fig. 4A-B). To test whether this compaction behavior is a general feature of PAR-binding proteins, we also tested the RNF146 WWE domain (36), which has a similar binding affinity to PAR but lower pI of 7.9 (Fig. 4A-C and S4). Upon binding to increasing concentrations of WWE, we did not observe any change in FRET signals and hence persistence length. These observations indicate that not all PAR-binding proteins induce conformational compaction of PAR, and that the presence of intrinsically disordered together with positively charged regions may be a critical requirement for PAR compaction.

**Figure 4.**
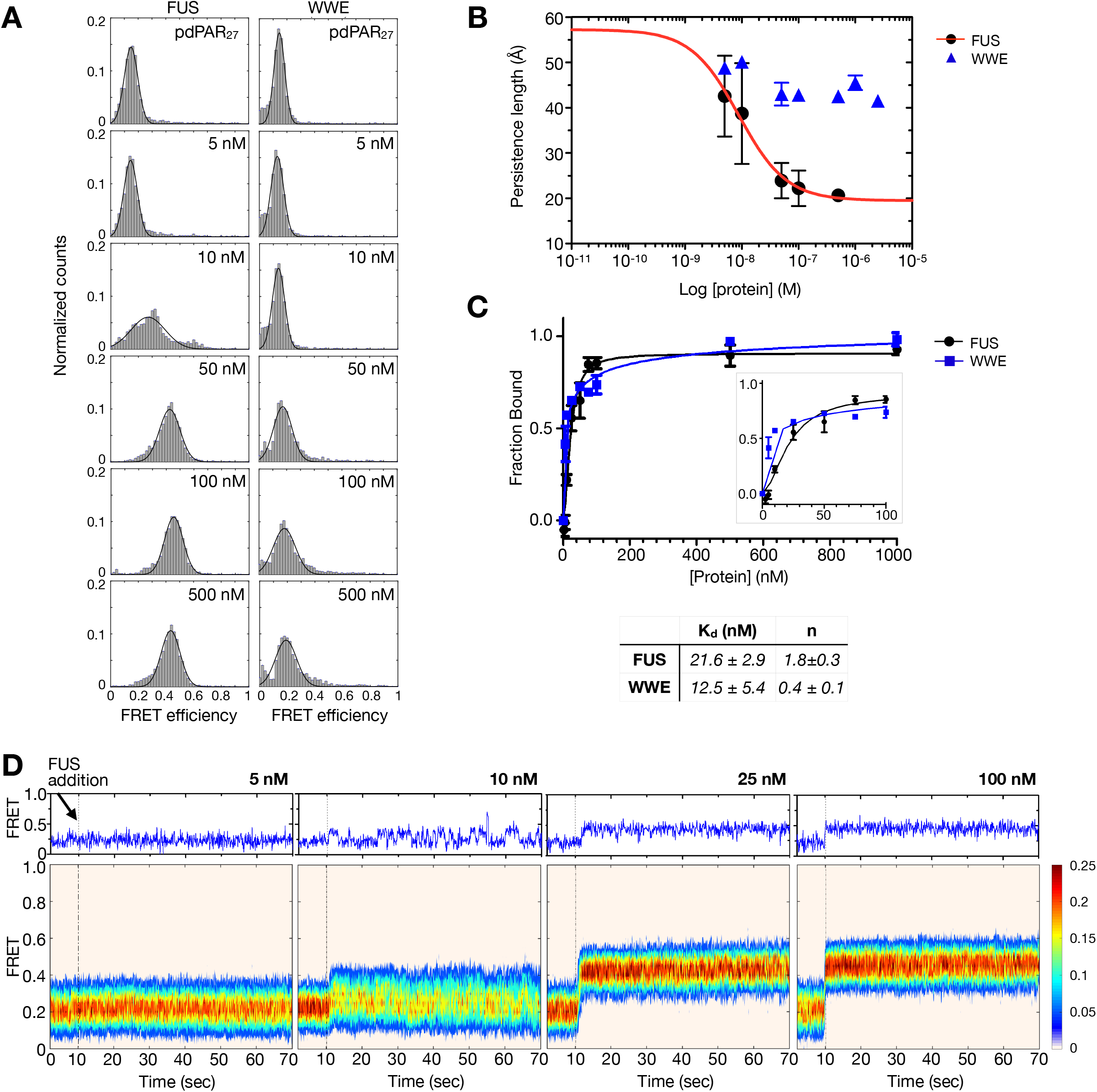
FUS and RNF146 WWE display different degrees of PAR compaction upon binding. **A.** FRET histogram of pdPAR27 with various concentrations of FUS and WWE. Single-peak Gaussian distributions were used to fit the histograms after removing donor-only population. The fit is shown as a solid black line, and the number indicate corresponding protein concentrations. **B.** The fractions of persistence length ***f*** of PAR were plotted over concentrations of WWE and FUS in log scale based on pdPAR27. Data were fitted in a power-law equation. **C.** The fraction bound was plotted over protein concentrations from EMSA measurement. The 0-100 nM range is zoomed out in a box for better visualization. The solid line is the fit to a modified Hill equation, with apparent Kd and Hill coefficient n indicated in the table. **D.** Real-time fluorescent measurement of pdPAR_27_ upon the addition of FUS. The dotted lines mark the introduction of FUS (flow at ~10 s) at indicated concentrations. The heat maps are generated by overlaying 80-120 time traces without synchronization of binding step. A representative time trace is shown on the top.

To further characterize the dynamics of FUS-mediated PAR compaction, we performed a real-time flow experiment in which smFRET videos were taken while adding different concentrations of protein (37). These real-time videos allowed us to capture the initial moment when FUS contacts PAR (Fig. 4D). We then compiled heatmaps using 80–120 video traces to analyze the pattern of FUS–PAR interaction. At 5 nM FUS, no appreciable change in the FRET efficiency was observed compared with the probe alone. The addition of 10 nM FUS induced an immediate increase in FRET efficiency with frequent fluctuations, indicating that PAR undergoes rapid conformational transitions, which may be explained by the transient compaction of PAR induced by FUS monomers in solution (34). Such rapid transitions are consistent with the lack of dominant patterns at a particular FRET efficiency in the heatmap due to the fluctuations of FRET values between 0.2 (unbound) and 0.5 (bound). In contrast, 25–100 nM FUS resulted in a dominant pattern in the heatmap, indicating a stable compaction of PAR upon multiple FUS binding at these concentrations. Consistent with this premise, the electrophoretic mobility shift assay (EMSA) also indicated a sharp transition of binding of FUS to PAR from 10 to 25 nM (Fig. 4C inset and S4A). At 25 nM, slightly more than half of PAR was bound to FUS with an additional complex formed at equilibrium (complex #2 in Fig. S4A). Taken together, PAR resists compaction until FUS concentration reaches a threshold, and the transition from extended to compact states occurs within a narrow range of FUS concentration, resembling the pattern observed for multivalent spermine and metal cations.

## Discussion

Here, we examined the flexibility of PAR homopolymers using our newly developed smFRET probes which allowed us to examine and quantify this fundamental property of PAR. Our analyses revealed that PAR is significantly less flexible than DNA and RNA at physiological sodium concentrations, which may have important implications for their differential ability to bind multivalently and phase separate (38). Yet, PAR undergoes a sharp transition from extended to compact state under physiological levels of divalent Mg^2+^ and Ca^2+^, as well as tetravalent spermine (39). The strong dependence on cation valency may arise from the unique chemical structure of PAR. The bonds linking phosphate and ribose are likely free to rotate in response to interactions with cations, as indicated by molecular dynamics simulations (40), whereas the bonds linking two consecutive ribose units between monomers contribute to the limited rotation within PAR (Fig. 2B). Therefore, the interplay between flexibility and stiffness likely drives the sharp transition from extended to compact PAR state, which we observed upon cation binding. We did not see the sharp transition for DNA or RNA, suggesting that this feature might be unique for PAR. The sharp transition is likely due to the strong repulsive nature of the two adjacent negative charges within the PAR monomer, which makes this molecule especially responsive to multivalent cations that can neutralize both charges simultaneously. Such a sharp response may bestow PAR with a switch-like bending property, consistent with its role as a signaling molecule.

An outstanding question related to PAR function is how this homopolymer that lacks well-defined structure achieves binding specificity. We observed that different ionic environments induce different degrees of compaction, even in the same unit length of PAR. Therefore, PAR may respond to changes in ionic environment, which would impact how PAR binds to proteins, and ultimately yield specific functional consequences. For example, PAR identified in the extracellular matrix is complexed with Ca^2+^ (29, 41). Our data revealed that PAR exists in the most compact states when Ca^2+^ concentration was in 1–4.5 mM range, which agrees with the previous observation that PAR condenses into large microscopically visible structures in similar Ca^2+^ concentrations (29). Therefore, cation binding drives structural reorganization of PAR, which may result in higher order structural features that contribute to specificity. Furthermore, a single PAR homopolymer can assume a range of conformations depending on cation concentration, suggesting additional layers of functional regulation dependent on the ionic environment.

In general, PAR-binding regions of proteins are structurally diverse, ranging from folded domains to intrinsically disordered sequence stretches (42, 43). Here, we showed that binding to different proteins impacts PAR flexibility differently. We observed that highly positively charged FUS protein that features intrinsically disordered regions effectively induces PAR compaction; however, a well-folded, slightly positively charged WWE domain does not induce compaction, despite being able to bind PAR equally well. Therefore, one may hypothesize that the ability of protein binding to result in PAR compaction depends on the flexibility of the protein and distribution of positive charge on the protein surface. In this hypothesis, the protein acts as a macromolecular polycationic platform that can neutralize PAR charge and accommodate conformational changes that accompany compaction. Interestingly, FUS protein may represent a special class of binding partners for PAR. Our recent studies indicate that PAR is a potent inducer of FUS condensation through transient interactions (34). We now reveal through the real-time single-molecule time traces that the transient FUS-PAR interaction results in brief spikes of FRET in PAR, indicating a highly transient compaction of PAR induced by the FUS contact. Our previous study showed that nearly all interactions with PAR are transient at low FUS concentration (≤ 10 nM), likely reflecting the initial stage of condensation. Below this concentration, PAR is in its most extended conformation and resists compaction. Within a narrow concentration range (10-25 nM) of FUS, PAR sharply switches to the compacted states, potentially altering the FUS conformation to condensation-proficient form. Future studies are warranted to investigate whether the changes in PAR mechanical properties contribute to FUS nucleation and condensation.

Overall, the smFRET probes we developed here allowed the study of PAR flexibility for the first time and may serve as a platform to systematically unravel the principles of PAR-driven recognition. Furthermore, our study indicates that homopolymeric PAR can exist in different conformations and compaction states. Although the nature of the structural changes will require future studies, including studies of PAR conjugated to target proteins as a model of PAR as a post-translational modification, our work suggests that a stiff homopolymer can gain specificity through regulated compaction.

### Limitation of the study

As a first approximation, we used the conventional WLC model to describe the flexibility of PAR. Our experimental data and simulated FRET efficiency indicate a strong agreement of fit with minimal RMSD, similar to single-stranded RNA and DNA (11, 13), suggesting that the WLC model provides a good approximation of PAR’s flexibility. However, we note that the persistence length based on WLC model alone may not completely capture the physics of the PAR chain at all salt concentrations, as is the case with other nucleic acids (44). Moreover, due to the finite distance sensitivity of FRET measurements in assessing end-to-end distance, more comprehensive investigations are warranted to systematically define the flexibility of PAR at varying lengths under different cation and protein conditions. These measurements should include the use of other biophysical techniques and the exploration of modified WLC (45–47), as well as non-WLC models (48).

## Materials and Methods

### Preparation of PAR

PAR was produced as a mixture and fractionated to defined lengths (49). Briefly, 20 mM NAD^+^, 0.1 mg/mL PARP5a catalytic domain, 2.5 mg/mL histones, 100 mM Tris buffer pH 8, 5 mM MgCl_2_, and 1 mM DTT in 0.5 mL reaction volume were incubated at 25°C for 1.5 h. After precipitation in 10% TCA, PAR was released by incubating the precipitate at 60°C for 1 h in 0.5 M KOH and 0.5 mM EDTA at 1000 RPM. The released PAR was adjusted to pH 9 with AAGE9 buffer (250 mM ammonium acetate pH 9, 6 M guanidium HCl, 10 mM EDTA). Then, the bulk PAR mixture was cleaned up using m-Aminophenylboronic acid–Agarose beads as described (49). 1 mL of cleaned-up PAR with absorption at 258nm = 75-125 measured on nanodrop was injected in DNApac-PA100 using mobile phase A (25 mM Tris buffer pH 9) and mobile phase B (25 mM Tris buffer pH 9 + 1M NaCl) and fractionated at 5 mL/min by the following gradient program: 6 min (0% B), 18 min (30% B), 30 min (40% B), 90 min (47% B), 150 min (49% B), 162 min (55%), 174 min (63% B), 174.1 min (100% B), 178 min (100% B). PAR of defined length was dried in SpeedVac or lyophilizer. 2-7 mer of PAR were desalted with Hi-Trap desalting column and FPLC. PAR 5-40 mer were desalted with centrifugal Amicon filter units (3 kDa cutoff). If needed, PAR was concentrated before storage in −20°C. Alkali release of PAR generates AMP on 1” termini, which was verified by mass spectrometry (Fig. S1D). Hereafter, amp-PAR_n_ will be referred to as PAR_n_. The PAR_n_ concentration was estimated based on the Beer-Lambert law A_258nm_=εLC using molar extinction coefficient (ε) of PAR (ε _PAR_ = ((number of ADP-ribose * ε _ADPr_ [13500 M^-1^.cm^-1^]) (50) + ε _dAMP_ [15400 M^-1^.cm^-1^]).

### Preparation of smFRET PAR Probe

The PAR probe was prepared in four steps.

#### Step 1 (Alkyne modification)

A 50 μL reaction containing 1 nmol of PAR at a defined length, 100 mM 1-methylimidazole-HCl buffer pH 7, 250 mM alkyne-amine linker (adjust pH 7-8 by HCl), and 100 mM EDC was incubated at 20°C overnight with shaking. The reaction was cleaned up using three different methods based on the size of PAR: ion pairing HPLC for 2-7 mer PAR, G25 micro spin for 8-9 mer PAR, and Monarch® RNA clean up kit following manufacturer’s instructions (specific for 15 nt RNA) for 10 mer PAR and above. After cleaning up, PAR was dried in SpeedVac.

#### Step 2 (Cy3 labeling)

A labeling reaction was performed at two scales (up to 0.5 nmol, 20 μL and 1 nmol, 50 μL) using OAS1:PAR:dATP-Cy3 in 1:10:20-50 molar ratio. For example, 1 nmol PAR, 40 μM dATP-Cy3 (Jena Bioscience # NU-835-CY3), 0.1 mg/mL polyIC, 2 μM OAS1 in 20 mM T ris, 20 mM Mg(OAc)_2_, 2.5 mM DTT pH 7.5 were incubated for 2 h at 750 RPM and 37°C. Then, polyIC was digested by adding RNase R, 20 mM Tris buffer pH 8, 100 mM KCl and incubated for another 1 h at 750 RPM and 37°C. The enzymes, digested polyIC, and unreacted dATP-Cy3 were removed by Monarch® RNA cleanup kit following manufacturer’s instructions for 15 nt-RNA. Any size below 10 mer was cleaned up by injecting into a C18 column using mobile phase A (100 mM TEAA pH 7.5) and mobile phase B (acetonitrile) at flow rate 1 mL/min and the following program: 0 min (5% B), 10 min (5% B), 40 min (60% B), 42 min (100% B), 45 min (100% B). Each fraction with absorption at 258 nm and 550 nm was collected and dried in SpeedVac. Alkyne-PAR-Cy3 concentration was estimated using molar extinction coefficient (ε) of fluorophore (ε _Cy3_ = 150000 M^-1^.cm^-1^) or PAR (ε _PAR_ = ((number of ADP-ribose * ε _ADPr_ [13500 M^-1^.cm^-1^]) + ε _dAMP-Cy3_ [20300 M^-1^.cm^-1^]).

#### Step 3 (Click ligation of DNA to PAR)

A 10 μL reaction was prepared in a PCR tube by adding reagents in the following order: 2.5 μM Cy3-labeled PAR, 50 mM HEPES pH 7.5, 200 mM NaCl, 5 μM azide-DNA, 20% DMSO, 0.5/2 mM Cu/THPTA complex, 10 mM sodium. The solution was incubated at 37°C for 2 h. CuSO_4_ and THPTA were premixed in 200 mM NaCl and incubated for 10 min. Sodium ascorbate was made fresh prior to the addition. The reaction was stopped by adding EDTA to a final concentration of 10 mM and conducting ethanol precipitation. 100 μL was injected to Bio SAX column using mobile phase B (25 mM Tris buffer pH 9 + 1 M NaCl) and fractionated at 1 mL/min by the following gradient program: 0 min (30% B), 20 min (80% B), 20.1 min (100% B), 23 min (100% B). The 200-250 μL fraction containing DNA-PAR-Cy3 was precipitated by mixing 2 μL of 2 mg/mL glycogen, 20 μL of 3 M NaOAc pH 5 and 825 μL EtOH at −80°C overnight (minimum 1 hour) and spun down at max speed for 30 min. After supernatant removal, the pellet was washed with 150 μL of ice-cold 70% ethanol. Air dry and store at −20°C for storage. Resuspend in 2-4 μL of mQ water/TE buffer. DNA-PAR-Cy3 concentration was estimated using molar extinction coefficient (ε) of fluorophore (ε _Cy3_ = 150000 M^-1^.cm^-1^) or DNA-PAR (ε _DNA-PAR_ = ((number of ADP-ribose * ε _ADPr_ [13500 M^-1^.cm^-1^]) + ε _dAMP-Cy3_ [20300 M^-1^.cm^-1^] + ε _DNA_ [171700 M^-1^.cm^-1^]). ε _DNA-PAR_ was used for the concentration calculation needed for the annealing step.

Labeling and click ligation may be performed in any order. However, the yield of DNA-PAR-Cy3 increases significantly if the labeling reaction is performed first. Additionally, for very short (7-9 mer) or very long (32-35 mer) PAR, DNA-PAR-Cy3 can only be generated if the labeling reaction is performed first. Summary of the yield percentage of smFRET PAR constructs: pdPARy_7_ (11.2, 19.1), pdPAR_12_ (12.9, 31.7), pdPAR_17_ (19.0, 12.0), pdPAR_22_ (13.1, 30.3), pdPAR_27_ (18.3, 19.6), pdPAR_35_ (18.3, 89.2). The first number refers to the yield of click ligation after purification and the second number refers to the yield of labeling before annealing.

#### Step 4 (Annealing)

Partially complementary biotin-DNA-Cy5 (strand 2) and DNA-PAR-Cy3 (strand 1) were annealed in a PCR tube by mixing at a 1.2:1 molar ratio in 50 mM Tris buffer, pH 7.5, 100 mM KCl for a total of 5 μL and heated to 95°C for 2 min, followed by slow cooling (2°C per min) to room temperature. Annealing was verified by gel electrophoresis on 6% DNA retardation gel.

### Preparation of smFRET RNA Probe

The smFRET RNA probe was prepared as described (23).

### Preparation of FUS and WWE Domain

His-MBP-FUS was purified as described (23). Purified His-MBP-FUS was exchanged to 50 mM HEPES pH 7.4, 500 mM NaCl, 20% glycerol, 2 mM DTT, and flash frozen in single aliquots in −80°C. FLAG-WWE domain from RNF146 was purified as described (51) and stored at −80°C in 20 mM Tris buffer pH 7.4, 200 mM NaCl, 5% glycerol, 1 mM β-mercaptoethanol.

### smFRET Measurement

All the smFRET measurements were carried out using PolyEthylene Glycol (PEG) coated quartz slide as described (16, 52). The microfluidic sample chamber was prepared using thinly cut double-sided tape between a PEG-passivated quartz slide and a coverslip. The edges between the slide and the coverslip were sealed with quick-drying epoxy glue. smFRET probes were immobilized on the slides via biotin-neutravidin linkage by flowing 50 μL of the following solutions sequentially: 1.) neutravidin (50 μg/mL) in T25 (20 mM Tris buffer pH 7.4, 25 mM NaCl) buffer, 2.) T25 buffer wash, 3.) pdPAR/pdU (fresh dilution to 10-25 pM), and 4.) T25 wash. Before imaging, 50 μL of imaging buffer (20 mM Tris buffer pH 7.5, 0.5 w/v % glucose, 1 mg/mL glucose oxidase, 4 μg/mL catalase, and 5-10 mM trolox-NaOH pH 8) supplemented with the appropriate amount of each ion (NaCl, MgCl_2_, CaCl_2_, and spermine hydrochloride) was applied to the surface. Imaging buffer for FUS-PAR and WWE-PAR interactions were supplemented with 100 mM and 72 mM KCl, respectively, and the appropriate proteins.

### Data Acquisition and Analysis

All smFRET experiments were performed at room temperature using a custom-built prism-type total internal reflection inverted fluorescence microscope (Olympus IX 71; Tokyo, Japan), equipped with a solid-state 532 nm and 634 nm diode laser (Compass315M; Coherent, Santa Clara, CA) to excite Cy3 and Cy5, respectively as described earlier (16, 37, 52). The fluorescence from Cy3 (donor) and Cy5 (acceptor) were simultaneously collected using a water immersion objective (60X; Olympus). To separate the donor and acceptor signals, a dichroic mirror (cutoff = 630 nm) was used, and finally signals were projected on an EMCCD camera (iXon 897; Andor Technology, Belfast, UK) side-by-side. All the data were recorded with 100 ms frame integration time, unless otherwise stated. One field of view captures 75-300 molecules on average. The recorded data were processed using smCamera (16) and analyzed by customized MATLAB scripts (The MathWorks; Natick, MA). More than 2000 molecules (21 frames of 30 short movies) were recorded over different regions of the imaging surface to generate a FRET histogram for each experimental condition. The donor leakage was corrected based on the FRET-value of the donor-only molecules. Further, to exclude the donor-only contribution to the histogram at the low-FRET region, Cy3 and Cy5 molecules were excited sequentially (10 frames for Cy3, 1 frame dark, and 10 frames for Cy5) by using the green and red lasers, respectively. Each FRET histogram was normalized and fit to a Gaussian distribution using the “Gauss1” model in MATLAB.

For real-time measurements, a plastic reservoir was adhered above the inlet hole at one end of the chamber using the top of a pipet tip. The corresponding outlet hole at the opposite end of the chamber was connected to a syringe pump (Harvard Apparatus, MA, USA) via a silicone tube. Protein in imaging buffer was loaded into the reservoir, and real-time FRET images were collected by pulling the solution through the chamber at a flow rate of 20 μL/s. Heatmaps of the flow experiments were constructed in MATLAB without synchronization.

### Electrophoretic Mobility Shift Assay (EMSA) and Quantification

Cy5-PAR_27_ was prepared as described in Step 2. Its interaction with His-MBP-FUS and WWE domain was examined in 6% native gel (acrylamide/bisacrylamide 37.5:1). 10 μL of His-MBP-FUS binding mixture was prepared by combining 0-1 μM FUS, 2 nM Cy5-PAR_27_ in 50 mM Tris buffer pH 7.4, 100 mM KCl, 2 mM MgCl_2_, 0.1 mg/mL BSA, and 10 mM β-mercaptoethanol. 10 μL of WWE domain binding mixture was prepared by combining 0-2.5 μM WWE, 2 nM Cy5-PAR_27_, 7.5 mM Tris buffer pH 7.4, 70 mM KCl, 4 mM DTT, 0.1 mg/mL BSA, and 0.1% Triton X-100. Both mixtures were incubated at 24°C for 30 min and loaded on 6% native gel (acrylamide/bisacrylamide 37.5:1) after mixing with 1 μL of loading dye (100 mM Tris buffer pH 7.4, 50% glycerol, 0.1 % wt/vol orange G). Gels were imaged using the fluorescence mode on a Typhoon 9200 scanner. Gels were visualized and quantified using FIJI software (53). The intensities of each band after background subtraction were used to determine the unbound fraction. The bound fraction (1-unbound fraction) was plotted on a logarithmic scale of various concentrations of protein. The data points were fit to a modified Hill equation (54) in Prism GraphPad (Version 5.01) to get the apparent K_d_.

### Measurement of b_0_ for PAR and polyU

The 3D structure of proteins bound to poly (ADP-ribose) or ADP-ribose, and polyU or ssRNA containing a stretch of uridines were visualized in PyMOL (The PyMOL Molecular Graphics System, Version 1.8.6.2 Schrödinger, LLC.). 4L2H, 5A7R, 6B09, 3V3L, 5W6X, 4B1H, 3X0L, 4IQY, 4RMJ, 3V2B, and 6D36 were used for PAR (7– 9, 36, 55–59). 3ND3, 3PU4, 5GMG, 5N94, 5LTA, 3ICE, 5AOR, 6JC3, 6R7G, 6HXX, and 6TZ2 were used for polyU (60–70). The shortest distance between two phosphorus or oxygen atoms within a monomer or adjacent monomer in three-dimensional space was measured in Angstroms. The mean and standard error of these values were calculated in Prism GraphPad. Then, the data were plotted using the ggplot2 package in RStudio (71).

### Fitting smFRET Data to the WLC Model

The FRET efficiency of pdPAR_n_ (n = 7, 12, 17, 22, 27, 35) and pdU_n_, (n = 20, 30, 40, 50, 70) in NaCl (25, 50, 100, 200, 400, 800, 1000, and 2000 mM) were obtained experimentally in 2-4 replicates. The experimental FRET data was fit to the WLC model as described (11) using a customized MATLAB script. We verified our script by recalculating δ, Lp, RMSD values using published FRET value of polyT (11). We were able to generate the same Lp and δ as reported (Fig. S2E and G).

The constant parameters in the MATLAB script are the following: monomeric length (*b_0_*), random walk step size (*δ*), Forster Radius (R_0_), and polymer length (range). The R_0_ of Cy3-PAR and Cy3-polyU was assumed to be the same as Cy3-DNA at 59 Å. Other parameters are as follows: *b_0_* =11.6 Å, *δ* = 13.8 Å, and range = 2-45 mer for pdPAR_n_ and *b_0_* = 5.9 Å, *δ* = 9.5 Å, R_0_ = 59 Å, and range = 5-70 mer for pdU_n_. The values of persistence length that resulted in the simulated FRET values with the smallest root-mean-squared deviation 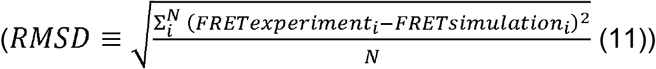 between the experimental and simulated FRET values were reported in Fig. S2G. The persistence length was calculated for each NaCl concentration. The simulated and experimental FRET values were exported to Prism GraphPad for visualization.

To obtain the step size (d) value, the above process was performed for d = 1-25 Å, with an increment of 0.5-1 Å. The RMSD values were then averaged over all NaCl concentrations and plotted over step size (Fig. S2D). The data was fit to a third-order polynomial equation, and the minimum value was used as the constant value for step size in the simulations described above.

To obtain the contour length, n * *b_0_* was used in the above simulation. This calculation of contour length does not consider the contribution of AMP (result of PAR preparation) and dAMP (result of labeling) on both termini of PAR. Thus, further simulations were performed by adding the length of AMP and dAMP to the contour length. Significant differences in persistence length or improvement in fit to experimental values were not observed.

### Data and Code Availability

The customized MATLAB scripts are deposited in GitHub (https://github.com/aLeungLab/WLC_FRET_Simulation) and the main experimental data are available in SI Appendix, Supplementary Tables. smCamera can be downloaded for free from the Ha lab (https://ha.med.jhmi.edu/resources). The MATLAB code for Monte Carlo computation was a generous gift from the Ha lab. In order to automate all calculations for PAR, polyU, and polyT, two patches were added to the Monte Carlo MATLAB script. The first patch allowed for input of the constant and variable parameters. The second patch added the minimization function to find the optimal parameters resulting in the best fit to experimental FRET values.

## Acknowledgement

We thank Drs. Phil Sharp, Sarah Woodson, Lois Pollack, and the members of the Leung lab for comments on the manuscript. We thank Dr. Taekjip Ha for MatLab scripts and discussion to analyze the FRET data in calculating persistence length. We acknowledge funding support for this work from the National Institutes of Health grants T32-CA009110 (M.B.) and RF1-AG071326 (A.K.L.L. and S.M.). The ELTA technology, which enabled the development of the smFRET probe, was funded by R01-GM104135 (A.K.L.L.).

## Supporting information

**Figure S1.**
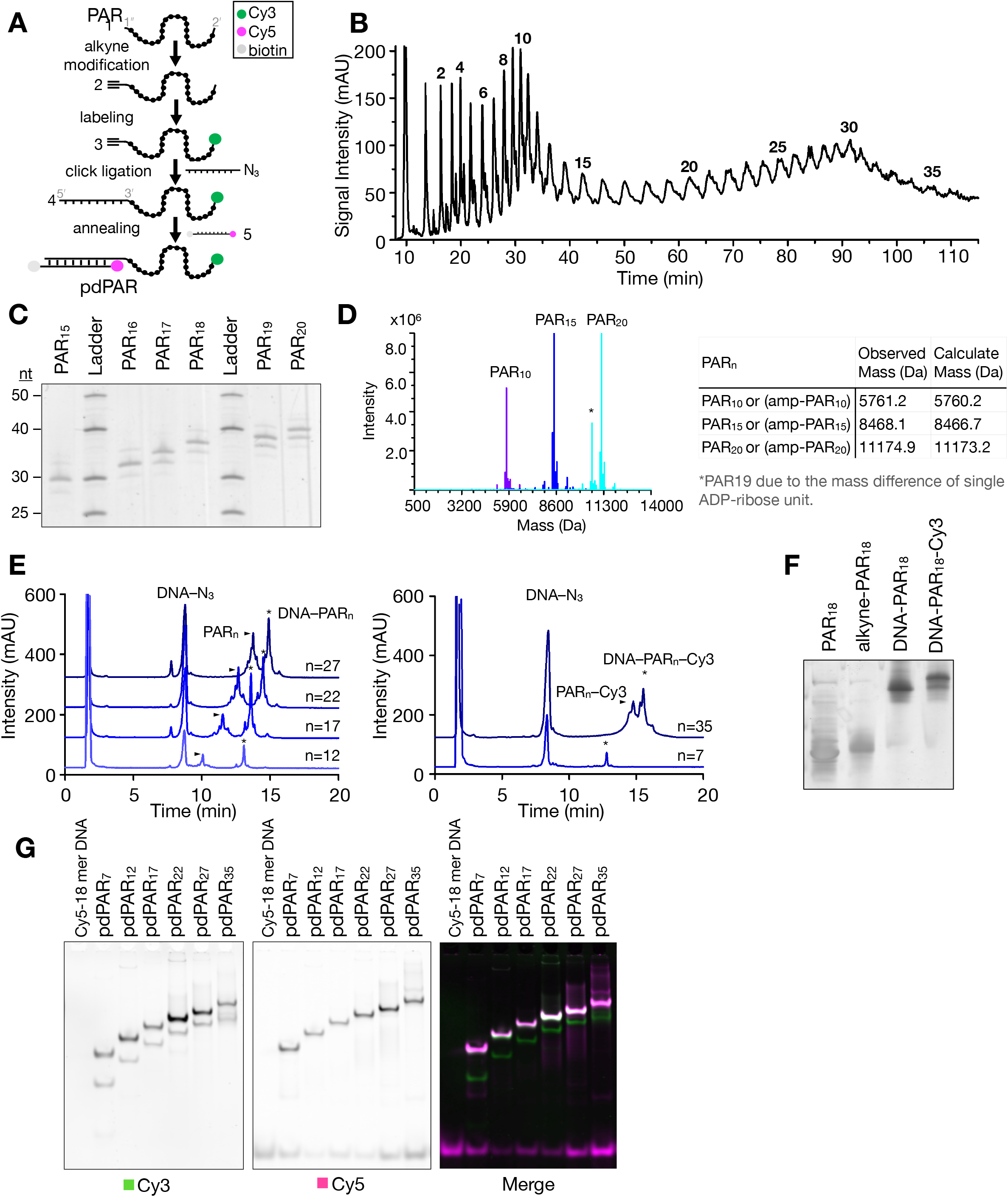
Development of smFRET probe to measure PAR flexibility (related to Fig.1). **A.** Synthetic steps to prepare pdPAR constructs of various lengths. **B.** HPLC chromatogram at 258 nm of PAR resolved to defined lengths of PAR. Peak number indicates the number of monomeric units. **C.** 20% urea-PAGE analysis of PAR_15-20_ after HPLC fractionation as compared to ssDNA ladder. The gel was stained with SYBR gold. **D.** Deconvoluted ESI-MS spectra of 10 mer, 15 mer, 20 mer PAR, with the observed and calculated mass indicated. **E.** HPLC chromatogram at 258 nm showing the isolation of reactant (DNA-N_3_ and PAR_n_) from chimeric DNA-PAR after step 3 in panel A. **F.** 15% urea-PAGE analysis of PAR_18_ and intermediates needed for the preparation of chimeric DNA-PAR_18_-Cy3. **G.** 15% PAGE native gel analysis of the final probe pdPAR_7-35_, indicated with Cy3 (green) and Cy5 (magenta) signals.

**Figure S2.**
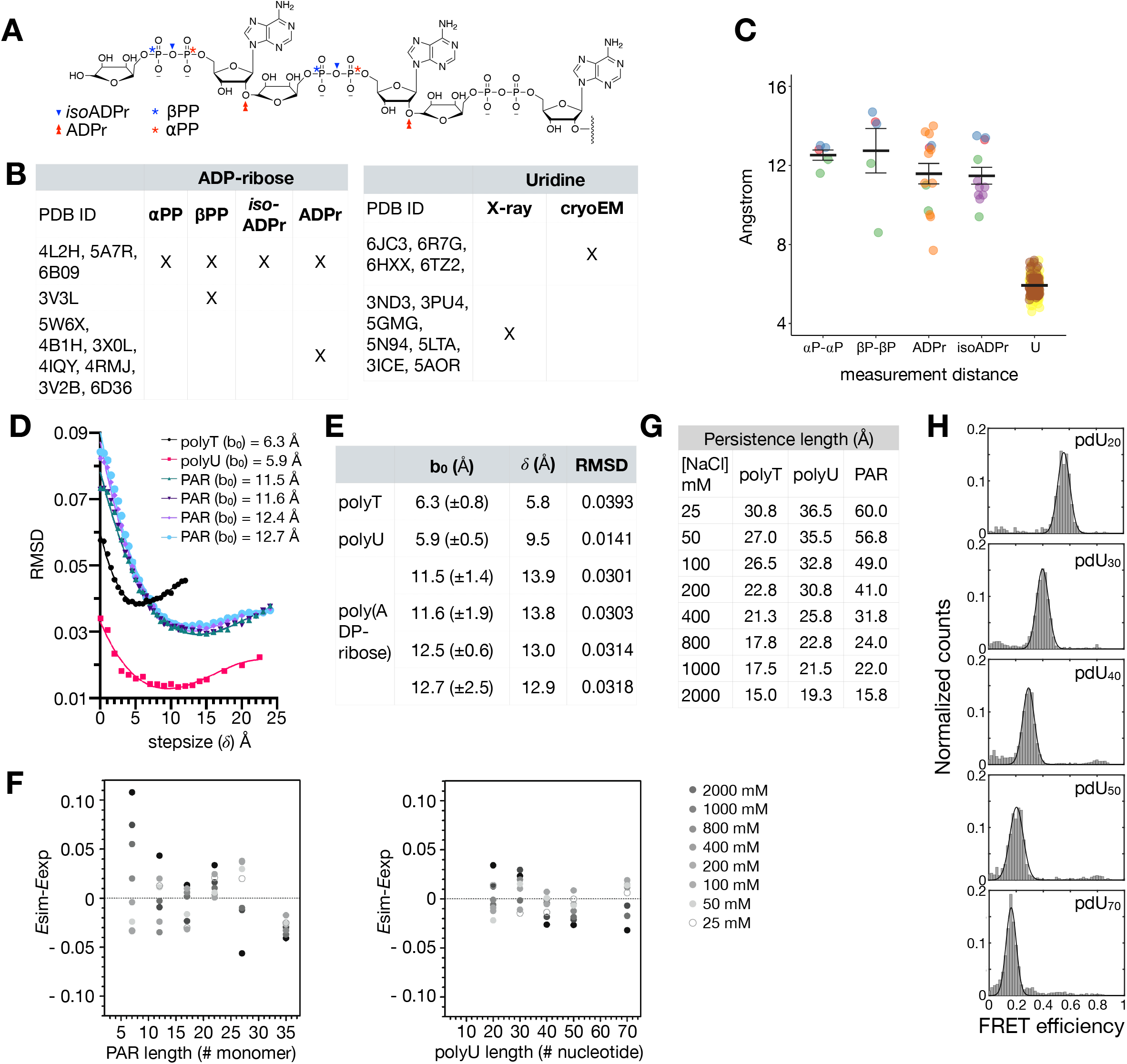
Worm-like chain model parameters and supporting calculation. **A**. Chemical structure of PAR, with marks indicating which substructures were used for bo measurements. **B.** PDB ID of ADP-ribose/PAR- or polyU-bound protein structures used for bo measurements. **C.** Scatter column plot of four b_0(PAR)_ and b_0(polyU)_. For PAR, colors indicate each measurement based on the structure with the same PDB. For polyU, colors indicate the measurement based on either X-ray crystallography or cryo-electron microscopy. **D.** Optimization of stepsize (*δ*), a parameter of biased random walk needed for energy transfer simulation (***E***). Each color corresponds to b_0(polyU)_, four b_0(PAR)_, and b_0(polyT)_, with the associated lines fitted with a third-order polynomial function. **E.** The final step sizes and minimal values of each fitted line in panel D are tabulated based on the corresponding b_0_. The RMSD values indicate the deviation between simulated (*E*) and experimental (*E*_FRET_) energy transfer values for the corresponding constant value, b_0_ and stepsize (*δ*). For polyT, the b_0_ value was based on Ref. 11. **F.** Residual plot indicates the deviation of experimental values (*E*_FRET_) from simulated values (*E*_FRET_). **G.** The persistence length of PAR_(b0=11.6 *δ*=13.β)_, polyU(b0=5.9 *δ*=9.5), and polyT(b0=6.3 *δ*=5.8) for NaCl from 25 mM to 2 M. For polyT, the values were recalculated based on FRET values from Table 1 in Ref. 11 and *δ* from panel E. **H.** FRET histogram of pdU_20-70_ at 200 mM NaCl. Single-peak Gaussian distributions was used to fit the histograms after removing donor-only population. The fit is shown as a solid black line. *Published values of poly T_40_ were used as comparison in panels D-F.

**Figure S3.**
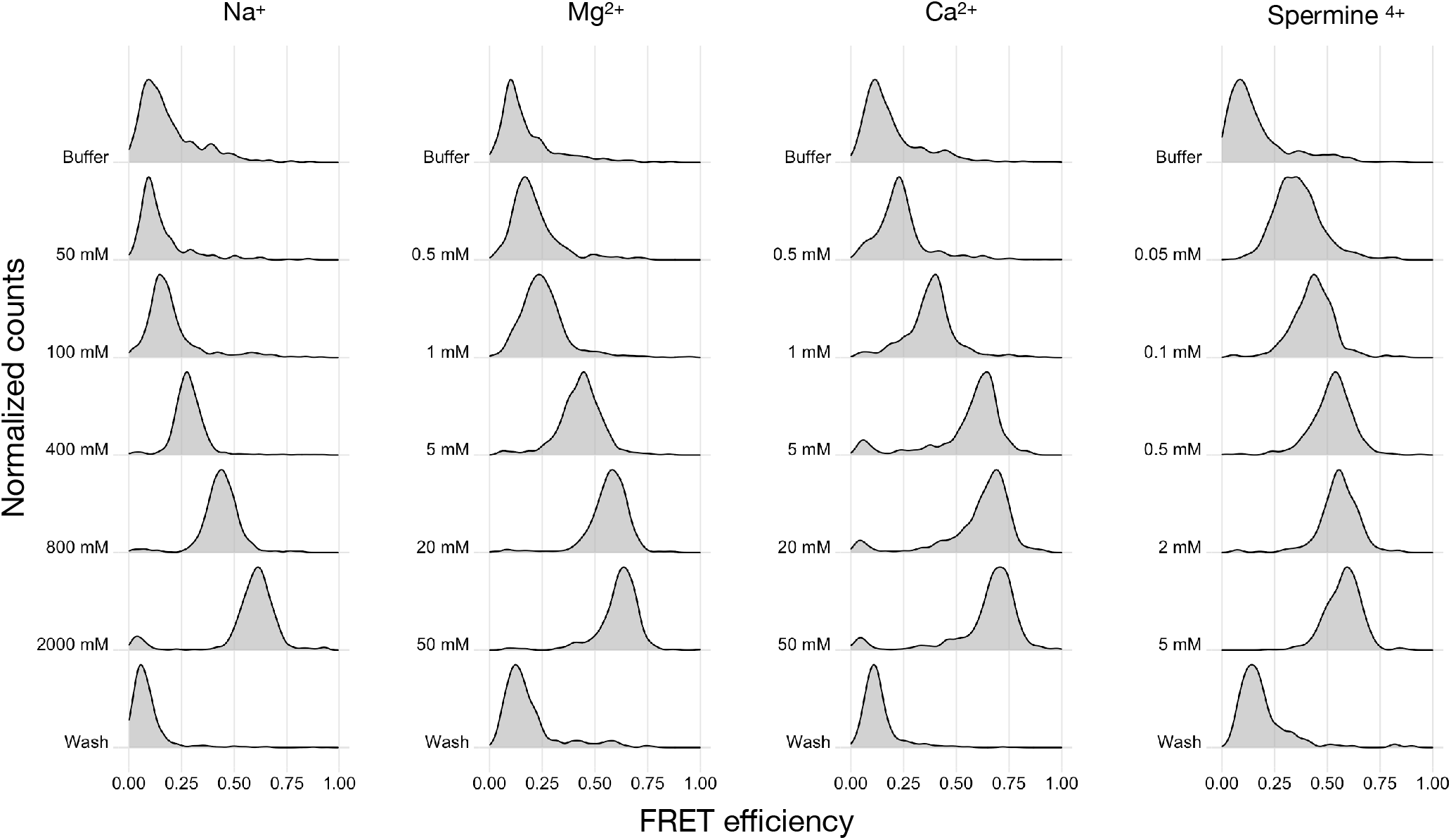
PAR is compacted by metals and spermine in a valency-dependent manner. FRET histogram of pdPAR_27_ titrated in various cations of different valency.

**Figure S4.**
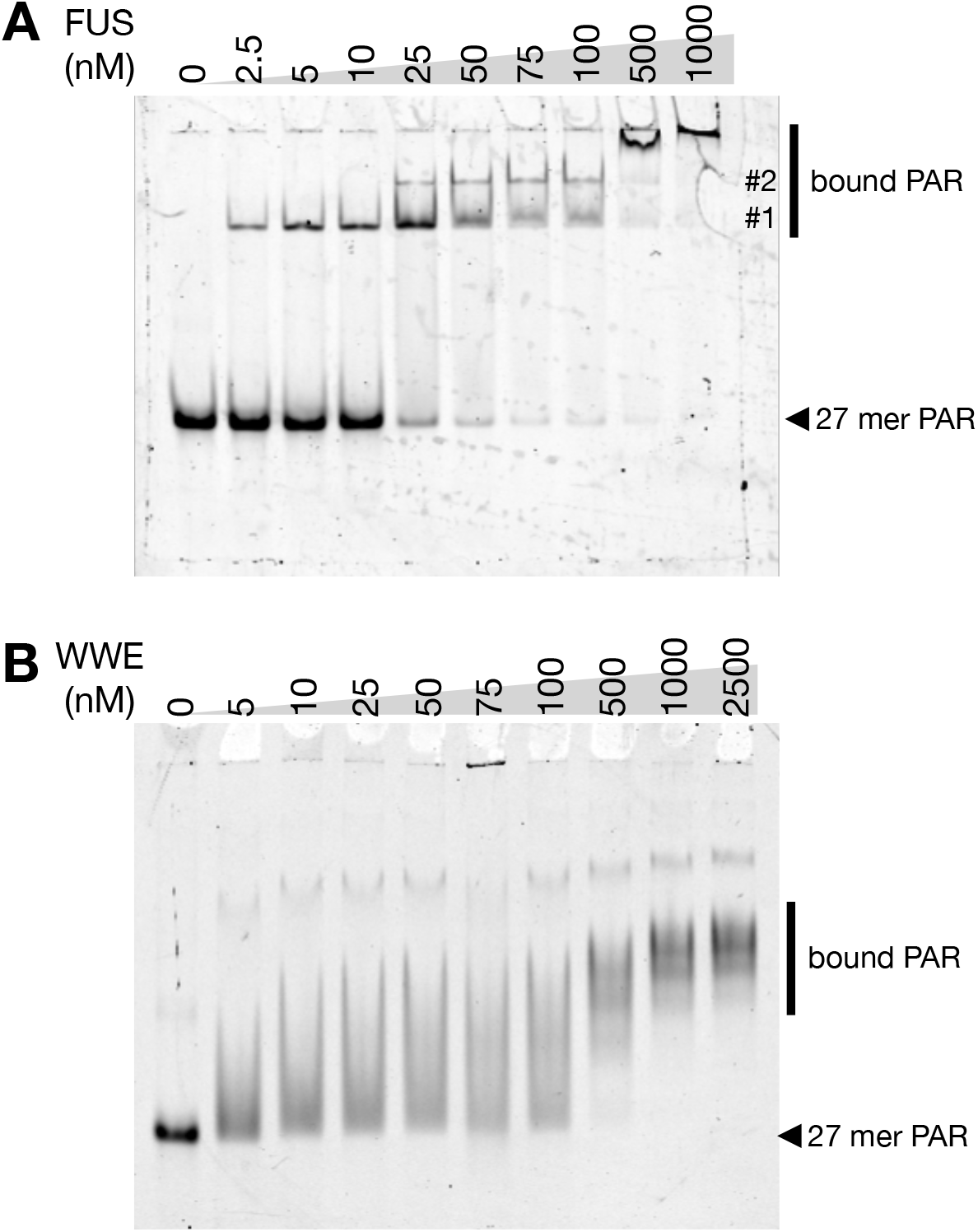
FUS and WWE domain display a similar affinity to PAR. 6% native PAGE gel analyses of the free and protein-bound of Cy5-PAR_27_ upon binding to **(A)** FUS or **(B)** WWE. The gels are visualized by Cy5 signal and #1 and #2 represent complex 1 and 2.

**Table S1.** smFRET efficiency for PAR and polyU of different lengths.

**A**
| [NaCl]<br>mM | pdPAR<sub>7</sub> | pdPAR<sub>12</sub> | pdPAR<sub>17</sub> | pdPAR<sub>22</sub> | pdPAR<sub>27</sub> | pdPAR<sub>35</sub> |
| --- | --- | --- | --- | --- | --- | --- |
| 25 | 0.55±0.00 | 0.22±0.08 | 0.19±0.02 | 0.10±0.02 | 0.07±0.03 | 0.10±0.01 |
| 50 | 0.56±0.01 | 0.24±0.08 | 0.19±0.04 | 0.13±0.00 | 0.08±0.02 | 0.11±0.02 |
| 100 | 0.64±0.00 | 0.30±0.08 | 0.27±0.02 | 0.15±0.00 | 0.11±0.04 | 0.12±0.01 |
| 200 | 0.69±0.01 | 0.44±0.04 | 0.28±0.07 | 0.23±0.01 | 0.15±0.04 | 0.17±0.01 |
| 400 | 0.75±0.01 | 0.58±0.03 | 0.41±0.05 | 0.33±0.01 | 0.24±0.04 | 0.25±0.01 |
| 800 | 0.80±0.00 | 0.68±0.06 | 0.55±0.03 | 0.44±0.01 | 0.39±0.05 | 0.34±0.00 |
| 1000 | 0.80±0.01 | 0.72±0.05 | 0.60±0.02 | 0.47±0.01 | 0.44±0.02 | 0.37±0.00 |
| 2000 | 0.82±0.03 | 0.78±0.05 | 0.71±0.03 | 0.60±0.03 | 0.61±0.04 | 0.51±0.00 |

| [NaCl]<br>mM | pdU<sub>20</sub> | pdU<sub>30</sub> | pdU<sub>40</sub> | pdU<sub>50</sub> | pdU<sub>70</sub> |
| --- | --- | --- | --- | --- | --- |
| 25 | 0.46±0.01 | 0.32±0.03 | 0.24±0.04 | 0.17±0.01 | 0.12±0.01 |
| 50 | 0.51±0.02 | 0.30±0.01 | 0.24±0.02 | 0.19±0.02 | 0.12±0.01 |
| 100 | 0.55±0.02 | 0.34±0.00 | 0.26±0.03 | 0.20±0.00 | 0.13±0.01 |
| 200 | 0.57±0.01 | 0.39±0.01 | 0.30±0.01 | 0.23±0.02 | 0.14±0.00 |
| 400 | 0.64±0.00 | 0.45±0.06 | 0.36±0.03 | 0.30±0.03 | 0.19±0.01 |
| 800 | 0.69±0.02 | 0.52±0.02 | 0.43±0.01 | 0.36±0.00 | 0.25±0.00 |
| 1000 | 0.71±0.00 | 0.53±0.01 | 0.46±0.01 | 0.39±0.00 | 0.26±0.00 |
| 2000 | 0.73±0.04 | 0.57±0.03 | 0.52±0.01 | 0.43±0.01 | 0.33±0.03 |

**B**
| [MgCl<sub>2</sub>]<br>mM | pdPAR<sub>27</sub> | pdU<sub>50</sub> |
| --- | --- | --- |
| 0.05 | 0.11±0.01 | 0.14±0.00 |
| 0.1 | 0.07±0.05 | 0.14±0.00 |
| 0.5 | 0.16±0.02 | 0.16±0.00 |
| 1 | 0.24±0.00 | 0.17±0.01 |
| 5 | 0.46±0.03 | 0.24±0.01 |
| 20 | 0.59±0.02 | 0.30±0.00 |
| 50 | 0.65±0.02 | 0.35±0.01 |

**Table S2.** PAR is compacted by metals and spermine in a valency-dependent manner. The concentration of cations at mid point of Lp from Fig. 3B (***C***_mid_) was extracted, and the range is presented as 95% confidence interval. ***n*** is exponent of the fitted equation (± std error). Experimental value of polyU50 as well as published values of polyU40 (Ref. 13) and poly T_40_ (Ref. 13) were used as comparison.

|  | PAR27 |  |
| --- | --- | --- |
| | $C_{\text{mid}}$ | $n$ |
| Na<sup>+</sup> (mM) | 404.5 (349.3-480.5) | 4.02 ± 0.51 |
| Mg<sup>2+</sup> (mM) | 1.0 (0.8-1.3) | 2.80 ± 0.50 |
| Ca<sup>2+</sup> (mM) | 0.6 (0.5-0.7) | 3.49 ± 0.72 |
| Sp<sup>4+</sup> (μM) | 24.0 (18.8-33.2) | 2.53 ± 0.32 |

|  | polyU50 |  | polyU40 |  | polyT40 |  |
| --- | --- | --- | --- | --- | --- | --- |
| | $C_{\text{mid}}$ | $n$ | $C_{\text{mid}}$ | $n$ | $C_{\text{mid}}$ | $n$ |
| Na<sup>+</sup> (mM) | 223.7 (198.6-251.8) | 1.86 ± 0.24 | 194.6 (182.0-209.2) | 1.62 ± 0.13 | 233.4 (220.0-248.4) | 2.01 ± 0.17 |
| Mg<sup>2+</sup> (mM) | 4.9 (4.2-5.8) | 1.62 ± 0.23 | 5.1 (4.6-5.6) | 1.43 ± 0.14 | 8.2 (7.5-9.2) | 1.52 ± 0.22 |

